# Influenza A virus hemagglutinin prevents extensive membrane damage upon dehydration

**DOI:** 10.1101/2021.11.22.469572

**Authors:** Maiara A. Iriarte-Alonso, Alexander M. Bittner, Salvatore Chiantia

## Abstract

While the molecular mechanisms of virus infectivity are rather well known, the detailed consequences of environmental factors on virus biophysical properties are poorly understood. Seasonal influenza outbreaks are usually connected to the low winter temperature, but also to the low relative air humidity. Indeed, transmission rates increase in cold regions during winter. While low temperature must slow degradation processes, the role of low humidity is not clear. We studied the effect of relative humidity on a model of influenza A H1N1 virus envelope, a supported lipid bilayer containing the surface glycoprotein hemagglutinin (HA), which is present in the viral envelope in very high density. For complete cycles of hydration, dehydration and rehydration, we evaluate the membrane properties in terms of structure and dynamics, which we assess by combining confocal fluorescence microscopy, raster image correlation spectroscopy, line-scan fluorescence correlation spectroscopy and atomic force microscopy. Our findings indicate that the presence of HA prevents macroscopic membrane damage after dehydration. Without HA, fast membrane disruption is followed by irreversible loss of lipid and protein mobility. Although our model is principally limited by the membrane composition, the macroscopic effects of HA under dehydration stress reveal new insights on the stability of the virus at low relative humidity.

## 1. Introduction

Seasonal influenza is an acute respiratory infection caused by influenza viruses (e.g., influenza A virus, IAV) circulating throughout the world and causing epidemics of the disease. Every year, up to 5 million cases of severe flu illness and 650000 flu-related deaths are reported [1]. Influenza activity follows largely predictable seasonal patterns, which are related to environmental conditions, such as temperature, ultraviolet light, and humidity [2] [3] [4]. Indeed, in temperate climates, the seasonal outbreaks occur mainly in winter, when humidity and temperature are low, while in tropical regions the patterns may be less pronounced. Among these conditions, the link between ambient humidity and virus transmission or survival has been subject of debate. Studies in animal models as ferrets [5] and guinea pigs [6] have demonstrated that cold temperatures and – independently – low relative humidity favor virus transmission. Epidemiological evidence suggests that humidity affects virus transmission by modulating virus survival [4], while other studies have associated also virus transmission and seasonality with either absolute humidity [7] or relative humidity [8]. Biophysical investigations have demonstrated that, at low humidity, the dehydration of respiratory droplets may affect the virus structure [9]. In stark contrast, recent research revealed the sustained infectivity and stability of IAV contained in human bronchial epithelial aerosols over a wide range of relative humidity [10].

Efforts to resolve the virus envelope structure at the molecular level have been based on the use of high-resolution techniques such as cryo-electron microscopy [11], cryo-electron tomography [12] or super-resolution fluorescence microscopy [13]. These studies have been mostly focused on revealing the spatial organization on the surface of the virus and usually have been carried out in solution or in ice (frozen buffer). Still, the detailed interactions between water and the molecular components of enveloped virus (specifically, in the case of low water amounts) have so far been neglected and are not completely understood.

In this study, we report unprecedented observations of the structural and dynamical changes taking place in models of the IAV H1N1 envelope after a complete cycle of hydration, dehydration and rehydration, by using a combination of fluorescence microscopy imaging, raster image correlation spectroscopy (RICS), line-scan fluorescence correlation spectroscopy (LSFCS) and atomic force microscopy (AFM).

Solid supported lipid bilayers (SLBs) have been extensively used as model membranes in biophysical studies due to their high stability and versatility in physiological conditions [14]. These phospholipid bilayers supported by solid substrates can be functionalized with proteins in a well-defined orientation and allow the study of membrane structure and dynamics [15] [16] [17] [18]. In this work, we present a combined AFM and fluorescence microscopy study of models of the IAV H1N1 virus envelope consisting of SLBs functionalized with the viral “spike” protein hemagglutinin (HA).

Fluorescence microscopy and AFM have been widely employed in virus research to perform molecular studies in physiological conditions [19] [20] [21]. The combination of these complementary techniques allows to understand complex membrane systems and characterize protein-protein, lipid-lipid and protein-lipid interactions [22]. In the context of fluorescence microscopy, advanced quantitative approaches such as RICS and LSFCS provide information on the dynamics of molecules by measuring e.g. local concentrations and diffusion coefficients of proteins and lipids, protein partitioning, or association and dissociation constants [23] [24]. LSFCS in particular minimizes photobleaching effects and provides more reliable results in dehydrated lipid samples, for which molecular diffusion is very slow [25]. The two-dimensional approach of RICS offers the exploration of the dynamics of molecules over a wide temporal and spatial scale, from microseconds between adjacent pixels in the x-direction, milliseconds between adjacent pixels in the y-direction, and seconds between successive images [26]. Finally, AFM provides structural information in native environments, from solution to variable humidity in air, or even vacuum. A detailed topography of the surface can be obtained from a three-dimensional scan of the surface with nanometric spatial resolution. This includes also adsorbed water layers, so that height variations can be correlated with air humidity. AFM studies on influenza viruses are not new [27] but have been usually performed only in liquid environment [28] [29] [30] or in desiccated state [31].

By combining all the above-mentioned techniques, we investigated here SLBs containing the IAV H1N1 surface glycoprotein HA and compared them against protein-free bilayers or bilayers containing a non-viral model protein (i.e., green fluorescent protein). We obtained detailed information on membrane dynamics and surface morphology at the macro and microscale -in solution and in air- under hydration, dehydration, partial rehydration and full rehydration conditions.

## 2. Materials and methods

### 2.1 Materials

1,2-Dioleoyl-*sn*-glycero-3-phosphocholine (DOPC), 1,2-dioleoyl-sn-glycero-3-[(N-(5-amino-1-carboxypentyl)iminodiacetic acid)succinyl] (nickel salt) (DGS-NTA-Ni), 1-palmitoyl-2-(dipyrrometheneboron difluoride)undecanoyl-sn-glycero-3-phosphocholine (TF-PC) and 1,2-dioleoyl-sn-glycero-3-phosphoethanolamine-N-(lissamine rhodamine B sulfonyl) (ammonium salt) (Rho-PE) were purchased from Avanti Polar Lipids (Alabaster, Alabama, USA) and used without further purification. Influenza A recombinant HA [A/Puerto Rico/8/1934 (H1N1)] was purchased from The Native Antigen Company (Kidlington, Oxford, United Kingdom). The protein was produced in mammalian HEK293 cells (≥ 95 % purity determined by SDS-PAGE from the manufacturer), providing the required post-translational modifications (i.e. glycosylation). The ectodomain protein fragment (Asp18-Gn529) was fused with 8 × Gly-Ser linker and a 6 × His-tag at the C-terminally located transmembrane domain. *Aequorea victoria* green fluorescent protein (GFP) was bought from Invitrogen (Carlsbad, California, USA). The recombinant protein was expressed in *Escherichia coli* (≥ 85 % purity determined by SDS-PAGE from the manufacturer) and was fused with 6× His-tag at the N-terminus. Alexa Fluor 568 succinimidyl ester (A568) and Rhodamine Red (Rho) were acquired from Thermo Fischer Scientific (Karlsruhe, Karlsruhe, Germany). Atto 488 amine fluorescent label, Dulbecco’s phosphate buffered saline (DPBS), absolute ethanol, acetone and silica gel microspheres were purchased from Sigma-Aldrich (Steinheim, Westphalia, Germany). The hydrophobic ink was obtained from Electron Microscopy Sciences (Hatfield, Pennsylvania, USA). Glass dishes (D35-20-0-TOP) were bought from Cellvis (Sunnyvale, California, USA) and the glue NOA63 was purchased from Norland (Cranbury, New Jersey, USA).

### 2.2 Fluorescence labeling

Influenza A HA was conjugated with the Alexa Fluor 568 succinimidyl ester dye. The protein was incubated overnight at 4 °C in a 1:1:0.1 molar ratio of protein, reactive dye and sodium bicarbonate buffer (1 M NaHCO_3_, pH 8.3–9.0). Free dye was removed by dialysis using a 12000 MWCO mini-dialyzer (Carl Roth GmbH & Co., Karlsruhe, Germany) against DPBS buffer, pH 7.4, overnight at 4 °C. Protein concentration and labeling efficiency were determined by absorbance at 280 nm and 578 nm, respectively. The HA concentration was estimated by using the molar extinction coefficient of the protein monomer (ca. 86000 M^−1^cm^−1^). Protein concentrations were generally between 4.5 and 7.5 μM, while labeling efficiencies were of 1 dye molecule every 4-10 monomers.

### 2.3 Preparation of SLBs

SLBs were prepared according to the vesicle fusion method [32]. For the formation of multilamellar vesicles (MLVs), 90 mol % DOPC and 10 mol % DGS-NTA-Ni were mixed in absolute ethanol and labeled either with TF-PC or Rho-PE. 0.25 mol % of fluorescent lipid was used for imaging and 0.01 mol % for quantitative fluorescence measurements. After solvent evaporation, the lipid film was rehydrated with DPBS to a final concentration of 4.1 mg/mL lipid. The MLV suspension was sonicated for 10 minutes at 40 °C to form small unilamellar vesicles (SUVs) and diluted 10-fold with the same buffer. 100 μL of suspension were deposited on a glass dish (AFM surface roughness (0.26 ± 0.02) nm) previously cleaned with absolute ethanol and milli-Q water. To perform AFM in solution, the liquid droplet was confined with a hydrophobic pen. For fluorescence microscopy and AFM in air, sample confinement was carried out by attaching a plastic cylinder of 7 mm diameter to the glass with a thin layer of NOA63 glue. Vesicle fusion and bilayer formation were induced by addition of 5 mM of calcium chloride (CaCl2) and unfused SUVs were removed by extensive rinsing with DPBS buffer (pH 7.4).

### 2.4 Protein binding to the SLBs

Labeled HA was added to the previously prepared SLBs to a final concentration of 2 μM. The proteins were incubated for 50 minutes to achieve high packing density on the SLBs and optimal binding time via Nickel (Ni2+)/hexahistidine-Tag (Ni/His-tag) interaction [33]. The incubation time was selected based on the time needed for the signal to reach a near-equilibrium level. Next, we obtained a binding curve using different HA concentrations, ranging from 0 to 1.75 μM. We estimated the maximum (saturation) concentration to be reached already at 1.75 μM and selected 2 μM as working concentration. For the experiments described in this work, the optimized binding parameters were used to prepare bilayers functionalized with HA (“HA-bilayers”) or His-tag GFP (“GFP-bilayers”).

### 2.5 Sample dehydration and rehydration

After the preparation of samples as described in 2.3 and 2.4 (referred to as “hydrated samples”), buffer salts were removed by rinsing with milli-Q water multiple times to avoid membrane defects due to the formation of crystals. For sample dehydration, ~ 60 % of the total water volume was pipetted out. The remaining water was removed for 48 h in a chamber with silica gel microspheres (20 ± 3) % relative humidity. These samples are referred to as “dehydrated samples”. Partial rehydration of SLBs was achieved by incubating the samples in a sealed chamber with (95 ± 3) % relative humidity for 24 h. The relative humidity of the chambers was continuously monitored with a mini data logger (174 H, Testo, Lenzkirch, Germany). These samples are referred to as “partially rehydrated samples”. To achieve full rehydration of the membranes (“fully rehydrated samples”), 150 μL DPBS buffer (pH 7.4) was added directly to partially rehydrated samples.

### 2.6 Confocal fluorescence microscopy

Confocal fluorescence microscopy imaging, RICS and LSFCS were performed on a LSM 780 Zeiss confocal microscope (Zeiss, Oberkochen, Germany) using a 40 × Apocromat 1.2 NA water immersion objective. Single color measurements were performed using an excitation at 488 nm provided by an Argon laser (for TF-PC or GFP). Excitation and emission light were split with a 488 nm dichroic mirror. Two-color measurements were performed with an additional excitation line at 561 nm provided by a laser diode (for the fluorophores Rho-PE, A568). In this case, a 488/561 nm dichroic mirror was used to separate excitation and emission. The microscope was calibrated before each experiment with Atto 488 (488 nm) and with Rho (561 nm) fluorescent dyes diffusing in solution. Typical counts per molecule were ca. 3.6 kHz for Atto 488 and 4.2 kHz for Rho. Daily variations were consistently within 10 % of these values.

Images of 512 × 512 pixels were acquired for hydrated samples using 488 nm laser excitation with ca. 4 to 8 μW (between 10 and 100 μW for 561 nm). For measurements in dehydrated or partially rehydrated samples, SLBs were sealed with a glass dish top and tape immediately after being removed from the incubation chamber. Manual scratching tests were performed on HA-bilayers after dehydration using a sharp metal tweezer as reported elsewhere [34]. Laser power for measurements in dehydrated or partially rehydrated SLBs varied accordingly to the type of sample, from 4 to 20 μW at 488 nm and from 10 to 100 μW at 561 nm. After full rehydration, 2 μL of a 1 mM TF-PC or Rho-PE solution were added directly on the protein-free bilayers and GFP-bilayers to improve imaging quality. This was not necessary for HA-bilayers as the fluorescence intensity remained high throughout the dehydration and rehydration cycle. All images were acquired with 3.15 μs, 12.61 μs or 25.09 μs pixel dwell time and processed with ZEN software version 3.0 (Carl Zeiss, Germany). The surface coverage of the dehydrated protein-free bilayers (number of images, n = 22), and the GFP-bilayers (n = 16) was calculated with ImageJ (ImageJ, Bethesda, Maryland, USA).

RICS measurements were performed in the framework of arbitrary-region RICS [35] [36]as previously described [23] [37]. Single frames of 256 × 256 pixels and 40 nm pixel size were acquired. Between 9 and 18 measurements were collected per sample in at least three replicates. Data in hydrated samples was acquired at 50.42 μs pixel dwell time, 0.42 μs line time and 7.75 s frame time. Measurements in dry or partially rehydrated samples were acquired at the lowest dwell time possible (177.32 μs pixel dwell time, 0.42 μs line time and 27.24 s frame time) and the laser power was adjusted to maximize the signal and minimize photobleaching for all species at each condition [26]. For 488 nm excitation, 5 μW (TF-PC) or 0.5-1 μW (GFP) excitation power were used in hydrated samples. 5-8 μW (TF-PC, GFP) were used for dehydrated and partially rehydrated samples. 5-8 μW (TF-PC) or 1 μW (GFP) were used for fully rehydrated samples. The A568 fluorophore used for protein labeling was always excited with ca. 1 μW laser power, while Rho-PE excitation was optimized based on the specific sample (5-10 μW excitation power for hydrated samples, 2 μW for dehydrated and partially rehydrated samples, and 2-20 μW for fully rehydrated samples). RICS data of images containing photon counts as a function of pixel position were analyzed using a custom-written MATLAB code (The MathWorks, Natick, MA). Diffusion results were plotted with GraphPad Prism version 8.0.1 for Windows (GraphPad Software, San Diego, California, USA) and the diffusion coefficients were expressed as the mean ± standard deviation (SD).

LSFCS measurements were performed as described in [38]. Briefly, a line scan of 256 × 1 pixels (pixel size 40 nm) was performed with 0.79 μs pixel scan time and 300000 lines were acquired for each measurement (141.82 s total time). The laser excitation power for 488 nm was 8 μW and the A568 dye used for protein labeling was excited with ca. 5 μW for all the tested conditions. Two samples were examined for each condition and at least three measurements were taken per sample. The LSFCS data (one channel per file) were exported as TIFF files, imported and analyzed in MATLAB as previously described [38].

### 2.7 Atomic force microscopy

Topography images were obtained on an Agilent 5500 microscope (Keysight, Santa Clara, USA) at (23 ± 1) °C, 0° scan angle and speed rates between 0.8 and 1.2 lines per second. The scanning was performed on at least a total of 7500 μm2 from different areas per sample, in three replicates per condition. The images were examined by using picoView 1.14 software (Keysight, Santa Clara, USA). Images of 256 × 256 pixels were acquired in hydrated samples in contact mode in DPBS buffer (pH 7.4). A DNPS silicon nitride probe < 10 nm nominal radius, 0.12 N/m spring constant and a resonant frequency of 34 kHz (Bruker, Madison, USA) was used for scanning. Prior to each measurement, the probe was cleaned with acetone and absolute ethanol. The liquid droplet over the SLBs had a diameter of ~ 20 mm and was confined using a hydrophobic ink. During the scanning, the sample was always covered with DPBS buffer and the set point was continuously adjusted to minimize the applied force. To obtain the dehydrated samples, we initially prepared the hydrated samples following the protocol described in 2.3 and 2.4 and afterwards dehydrated them as described in 2.5. For these samples, the solution was initially confined with a plastic cylinder (section 2.3) which was eventually carefully removed with tweezers. An environmental isolation chamber (Keysight, Santa Clara, USA) was mounted to the AFM, thus providing a complete sealed and isolated environment. A gentle and continuous stream of nitrogen was blown into this chamber till reaching (20 ± 3) % relative humidity. Scratch and scan tests were performed on HA-bilayers after dehydration. For this, a small area of the sample was completely removed with the AFM tip by scanning the sample in contact mode at high setpoint for three times. Afterwards, the tip was retracted from the surface and a larger area was scanned in acoustic mode (tapping mode). For partially rehydrated samples, a relative humidity of (95 ± 3) % was achieved by controlling the mixing ratio between a dry and a wet (water-saturated) stream of nitrogen. The sample was allowed to equilibrate overnight in the saturated chamber. The fully rehydrated samples were finally obtained by adding DPBS buffer (pH 7.4) to the partially rehydrated samples. Images of 512 × 512 pixels were obtained in acoustic mode for dehydrated, partially rehydrated and fully rehydrated samples. A Multi Al 75 probe (Budget sensors, Sofia, Bulgaria) with a nominal radius of < 10 nm, force constant of 3 N/m and resonance frequency of 75 kHz was used for imaging HA-bilayers and protein-free bilayers. Images on GFP-bilayers were acquired with a high resolution cantilever C14 Cr/Au, nominal radius of ~ 1 nm, force constant of 5 N/m and resonance frequency of 160 kHz (MikroMasch, Sofia, Bulgaria). AFM images were analyzed after scanning with Gwyddion software version 2.47 (Gwyddion, Czech Republic). All the topography images were first levelled by mean plane subtraction, aligned by height median and occasionally the z-excursions outliers were manually removed by e.g., correction of the scan line artefacts in the x-axis or misaligned segments within a single row. The surface coverage of the protein-free bilayers (number of images, n = 13), HA-bilayers (n = 19) and GFP-bilayers (n = 16) after dehydration was calculated with WSxM 5.0 software version 8.4 (Madrid, Spain). The height distribution histograms and height profiles were plotted with GraphPad Prism version 8.0.1 and the quantitative data was displayed as the mean ± SD.

### 2.8 Statistical Analysis

Statistical analysis was performed using GraphPad Prism version 8.0.1. Data sets were analyzed by unpaired t-tests (Mann-Whitney test) for comparing the same sample under different hydration conditions (e.g., partially rehydrated and fully rehydrated bilayers). In all the cases two tailed P-values were reported and the level of alpha was kept at 0.05.

## 3. Results

### 3.1 Fluorescence microscopy reveals protective effect of HA and irreversible loss of membrane fluidity after dehydration

In this study, we investigated membrane stability and dynamics in a IAV model system at different hydration conditions. We performed fluorescence microscopy imaging, RICS and LSFCS measurements in four conditions, namely 1) hydrated: in DPBS buffer immediately after preparation; 2) dehydrated: after removal of bulk water and desiccation at ~ 20 % relative humidity; 3) partially rehydrated: in equilibrium at ~ 95 % relative humidity and 4) fully rehydrated: obtained by adding DPBS buffer after 3).

IAV envelope models consisted here of SLB containing proteins at saturation surface density (Figure S1), to mimic the glycoprotein concentration found in actual virions [12]. We prepared membranes composed of DOPC 90 mol % with 10 mol % DGS-NTA-Ni (a nickel chelating lipid) and used trace amounts of either TF-PC or Rho-PE as fluorescent marker. As a model of IAV glycoproteins, we chose the HA ectodomain (Asp18-Gn529) of the strain H1N1 (A/Puerto Rico/8/1934) fused to a Gly-Ser linker and a His-tag at the C-terminus. The latter was used to couple to proteins to the SLBs, via interaction with DGS-NTA-Ni lipids [39]. We performed fluorescence imaging of the SLBs and quantified the final protein concentration by RICS. HA-bilayers were compared to GFP-bilayers, to assess whether membrane stability was affected by the protein’s identity or simply driven by e.g., protein crowding. Protein-free bilayers were used as negative control.

Initially, we performed fluorescence microscopy imaging and measured the diffusion of fluorescent lipids to investigate the overall bilayer integrity and the mobility of the SLBs components. We examined protein-free bilayers either with TF-PC or Rho-PE dyes and consecutively added the labeled HA or GFP, respectively. Figure 1 shows representative fluorescence microscopy images of SLBs labeled with either TF-PC (for protein-free- or HA-bilayers, A and B) or Rho-PE (for GFP-bilayers, C). Figure 2 shows the diffusion of fluorescent lipids quantified by RICS for the four tested levels of membrane hydration. As represented in Figure 1 A, we obtained homogenous protein-free bilayers after preparation in the hydrated state. After addition of HA (Figure 1 B) or GFP (Figure 1 C), the samples did not show macroscopic alterations, defects or aggregates.

**Figure 1.**
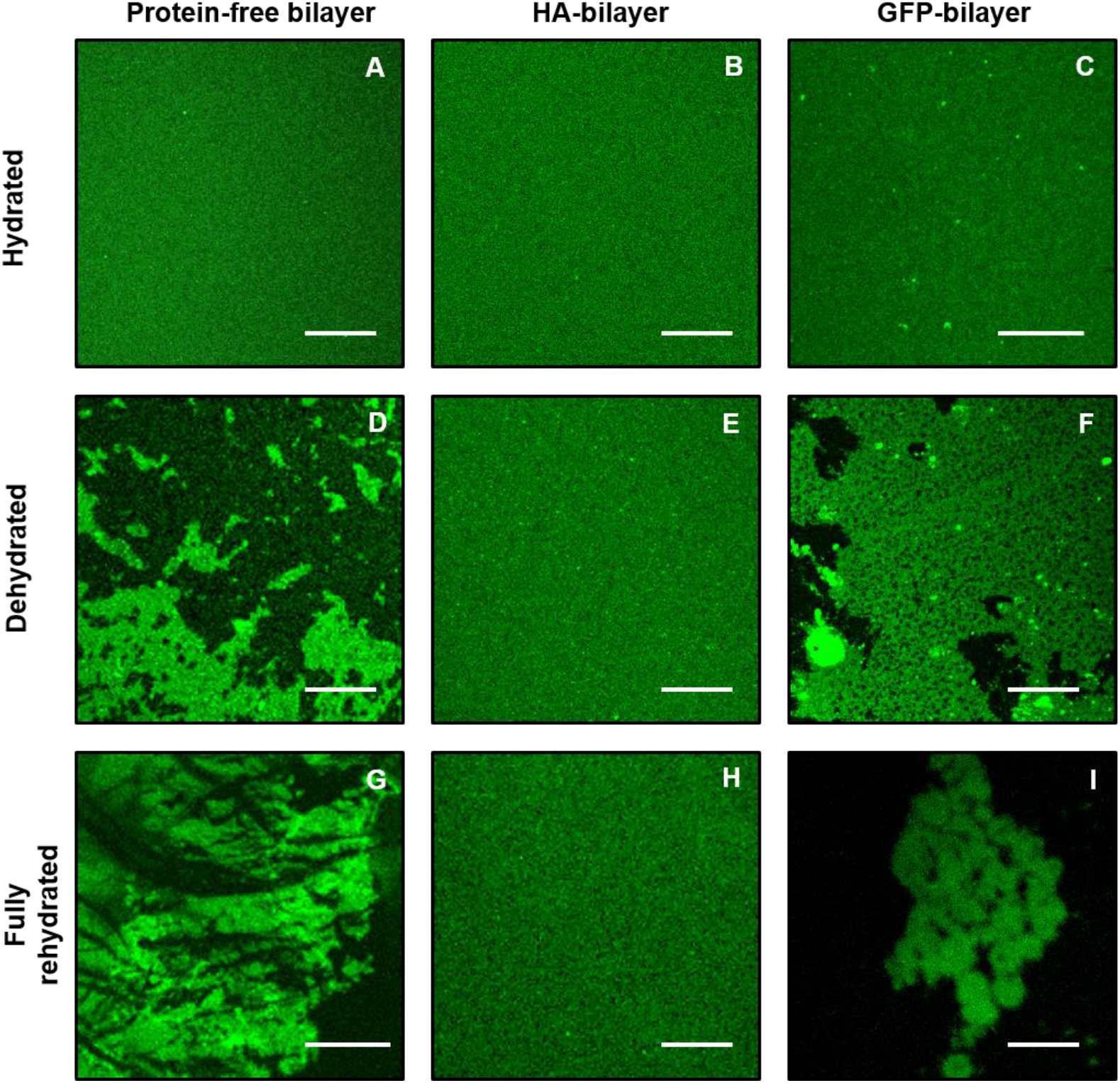
Fluorescence images of SLBs at different hydration conditions. (A-C) Representative images of hydrated SLBs in DPBS buffer immediately after preparation. (D-F) Representative images of dehydrated SLBs after removal of bulk water and desiccation at 20 % relative humidity. (G-I) Representative images of fully rehydrated SLBs in DPBS buffer. (A, D and G) Protein-free bilayers composed of DOPC 90 % DGS-NTA-Ni 10 % labeled with TF-PC. (B, E and H) TF-PC labeled bilayers composed of DOPC 90 % DGS-NTA-Ni 10 % conjugated to HA. (C, F and I) Rho-PE labeled bilayers composed of DOPC 90 % DGS-NTA-Ni 10 % conjugated to GFP. Fluorescence signal on G and I images were enhanced by adding 1 mM TF-PC or Rho-PE, respectively. Scale bars correspond to 20 μm.

**Figure 2.**
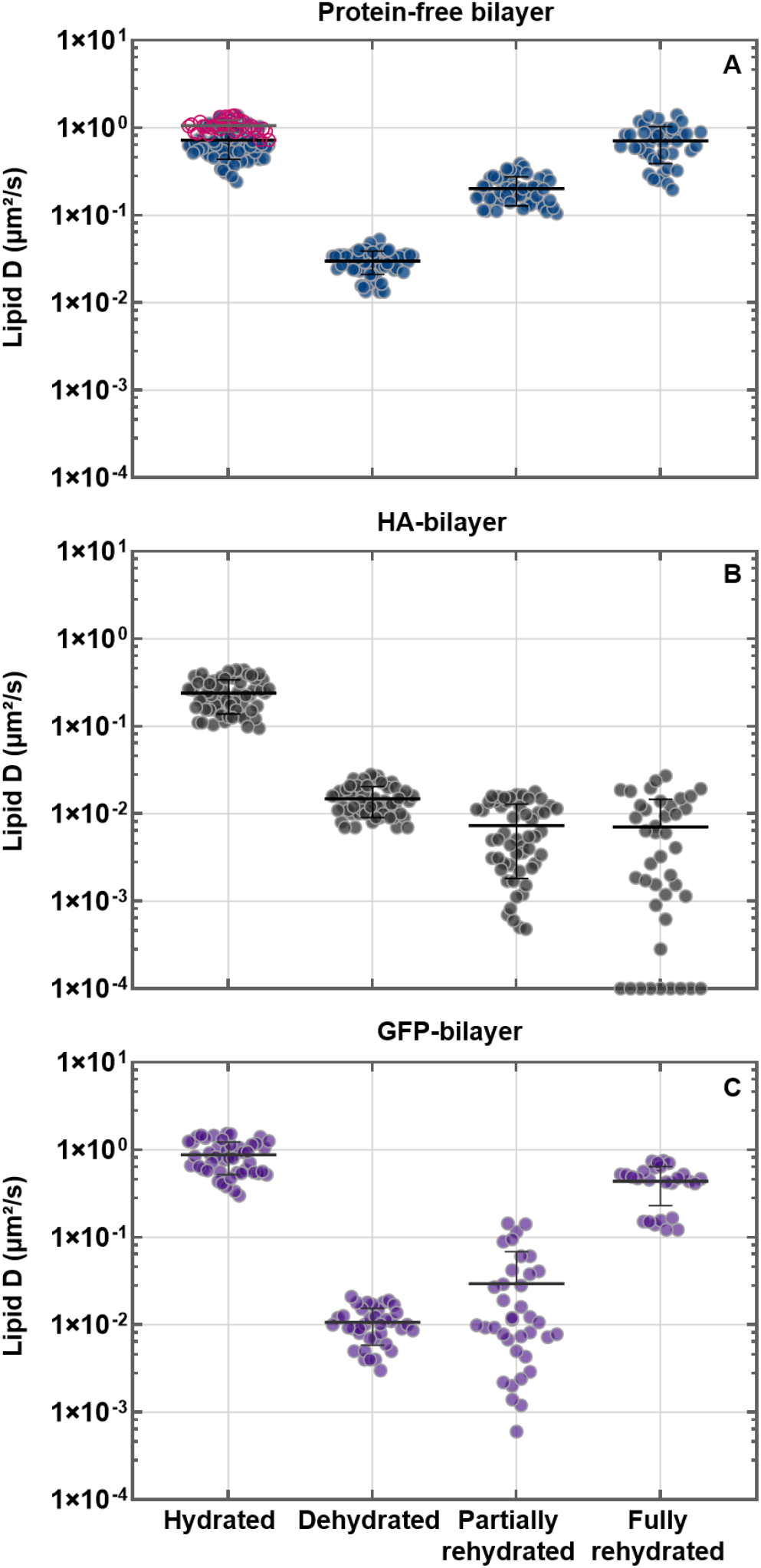
Diffusion coefficients (D) of lipids in SLBs measured via RICS at different hydration conditions. All examined SLBs contained DOPC 90 % and DGS-NTA-Ni 10 % and were labeled with 0.01 % TF-PC (A and B) or Rho-PE (C). Diffusion coefficients measured via RICS are shown for protein-free bilayers (A), bilayers containing HA (B), or bilayers containing GFP (C). Both proteins were anchored to the bilayer via His-tag and DGS-NTA-Ni. Note that the data shown in pink in (A) refer to the diffusion of Rho-PE. Data were obtained in at least three independent bilayer replicates (ca. 9-12 measurements per replicate). Individual points represent a single RICS measurement on one of the replicates. The horizontal lines show the mean D and the vertical lines the standard deviation. For better visualization, D values below 1 × 10^−4^ μm^2^/s were set as constant and equal to this minimum threshold.

The diffusion coefficients (D) of lipids (exemplified by the diffusion of the fluorescent lipid TF-PC) in protein-free bilayers was (0.7 ± 0.3) μm^2^/s (Figure 2 A “hydrated”). Once HA was added, the fluidity of the bilayer was reduced by ˃ 60 % ((0.2 ± 0.1) μm^2^/s, Figure 2 B “hydrated”). Rho-PE in protein-free bilayers shows a similar value, albeit slightly higher ((1.06 ± 0.17) μm^2^/s, Figure 2 A “hydrated”), compared to TF-PC. After binding of His-tag GFP to the SLBs, lipid mobility decreased by ca. 15 % ((0.9 ± 0.4) μm^2^/s, Figure 2 C “hydrated”).

Upon dehydration, the appearance of protein-free bilayers changed markedly with respect to the hydrated state. The samples showed extensive areas with structural damage, indicated by the presence of bare (i.e. devoid of lipids, dark) substrate regions (Figure 1 D). The membrane surface coverage decreased to (26.0 ± 17.0) %. On the other hand, HA-bilayers did not show any macroscopic defect and the fluorescence signal remained spatially homogeneous (Figure 1 E). This finding was further confirmed by a “manual scratching test”, for which a sharp steel tweezer was employed to remove the lipids, by applying a constant force within a selected region. The uncovering and the identification of the glass support serves to provide a defined (height or fluorescence signal) reference for further analysis. As presented in Figure 3 (A and B), within the scratched region we observed only background noise ((0.13 ± 0.04) A.U). Outside this region, the signal intensity was high ((2.41 ± 0.19) A.U) and comparable to what was previously observed (Figure 1 B and E). The presence of a bilayer in dehydrated samples was further confirmed by AFM imaging (Figure 3 C and D, section 3.4), showing a dip of ca. (3.8 ± 0.4) nm in correspondence of the scratched region.

**Figure 3.**
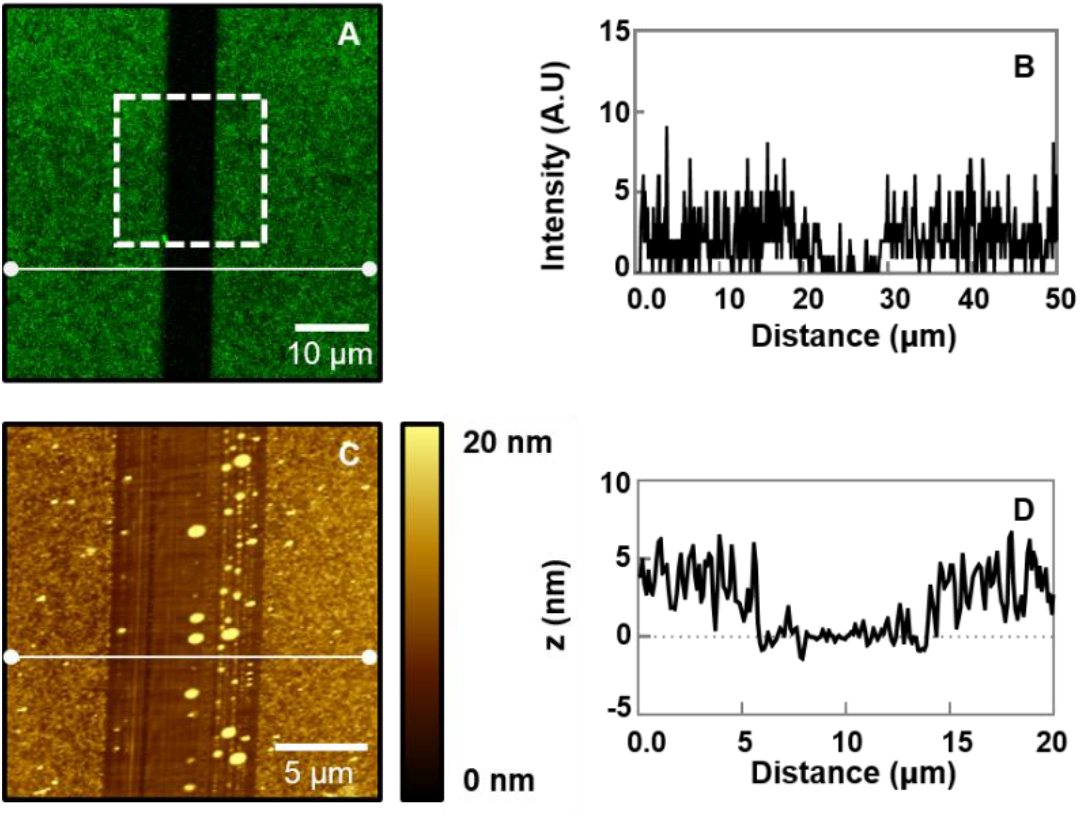
Manual scratching test performed on HA-bilayers after dehydration. (A) Representative fluorescence image of a scratched position on the DOPC 90 % DGS-NTA-Ni 10 % SLBs labeled with TF-PC and containing HA in the dehydrated condition (comparable to Figure 1 E). (B) Fluorescence intensity profile extracted from the horizontal line in (A). (C) Representative AFM topography image of the scratched sample (white square from (A)). (D) Height profile extracted from the horizontal line in (C).

Similarly to protein-free bilayers, GFP-bilayers presented macroscopic membrane disruption and exposure of the glass surface after drying (Figure 1 F). The surface coverage was reduced to (49 ± 26) %. Both protein-free bilayers and the GFP-bilayers exhibited high sample to sample variability (25.5 % SD and 17.0 % SD respectively).

RICS measurements were performed on random positions of the HA-bilayers. For GFP-bilayers and protein-free bilayers, measurements were performed specifically on non-damaged regions. Irrespective of membrane structure, we observed that the lipid mobility was greatly reduced after dehydration, as previously observed for protein-free bilayers [14]. In comparison to hydrated samples, lipids in protein-free bilayers displayed a D reduction of ~ 96 % ((0.03 ± 0.008) μm^2^/s, Figure 2 A “dehydrated”). Similarly, strongly reduced lipid dynamics were observed for HA-bilayers ((0.015 ± 0.006) μm^2^/s, Figure 2 B “dehydrated”) and GFP-bilayers ((0.011 ± 0.005) μm^2^/s, Figure 2 C “dehydrated”).

After partial rehydration, we could not detect significant large-scale structural changes of the SLBs by fluorescence microscopy (data not shown). RICS measurements were performed as described for the case of dehydrated samples. Diffusion within lipid patches in protein-free bilayers increased by ~ 30 % ((0.20 ± 0.07) μm^2^/s, Figure 2 A “partially rehydrated”). In contrast, lipid mobility on SLBs with HA or GFP remained low but exhibited a large data spread: (0.007 ± 0.005) μm^2^/s (Figure 2 B “partially rehydrated”) and (0.03 ± 0.04) μm^2^/s (Figure 2 C “partially rehydrated”) respectively.

Stronger differences were found after full rehydration in DPBS buffer (Figure 1 G-I). Microscopy images indicated an apparent increase in surface coverage for both protein free-bilayers and the GFP-bilayers (Figure 1 G and I, respectively). These bilayers were very inhomogeneous and bright structures could be found, possibly related to aggregates and lipids in non-bilayer phases. RICS measurements in specific points, where the bilayer was not damaged, indicated a diffusion recovery of ~ 99 % for the protein-free bilayers ((0.7 ± 0.3) μm^2^/s, Figure 2 A “fully rehydrated”), compared to hydrated samples. For GFP samples, lipid diffusion coefficients were ~ 50 % on those measured in the corresponding hydrated samples ((0.44 ± 0.21) μm^2^/s, Figure 2 C “fully rehydrated”). On the other hand, HA-bilayers appeared macroscopically homogenous, except for the presence of few aggregates (Figure 1 H). In these samples, no significant lipid diffusion could be observed ((0.007 ± 0.007) μm^2^/s, Figure 2 B “fully rehydrated”). Similar to the case of partially rehydrated samples, measurements exhibited in general a large variability.

### 3.2 RICS reveals a decrease of protein diffusive dynamics upon membrane dehydration

In order to obtain a more complete picture of the biophysical properties of the viral envelope models, we performed fluorescence microscopy imaging and analyzed the diffusion dynamics of proteins (i.e. membrane-anchored HA or GFP) within the examined SLBs (Figure 4). To this aim, a small fraction of proteins was labeled (section 2.2) and observed via fluorescence microscopy. RICS measurements were then performed to quantify the concentration and dynamics of the labeled proteins. From the fluorescence microscopy images in the hydrated condition we could observe homogenous protein adsorption on the SLBs both for HA and GFP (Figure 4 A and F). Taking into account the labeling efficiency, we could obtain the average total HA protein density on the bilayers, i.e. ~ 1.5 × 10^3^ HA monomers/μm^2^. Similar experiments were also performed for GFP-SLBs. Assuming that ca. 60 % of the anchored GFP is fluorescent [40] [41], RICS experiments indicated an average density of ~ 1.0 × 10^3^ GFP monomers/μm^2^ in our samples. Finally, diffusion coefficients were obtained for HA ((0.045 ± 0.026) μm^2^/s, Figure 4 D “hydrated”) and GFP ((0.05 ± 0.04) μm2/s, Figure 4 E “hydrated”) in the hydrated samples.

**Figure 4.**
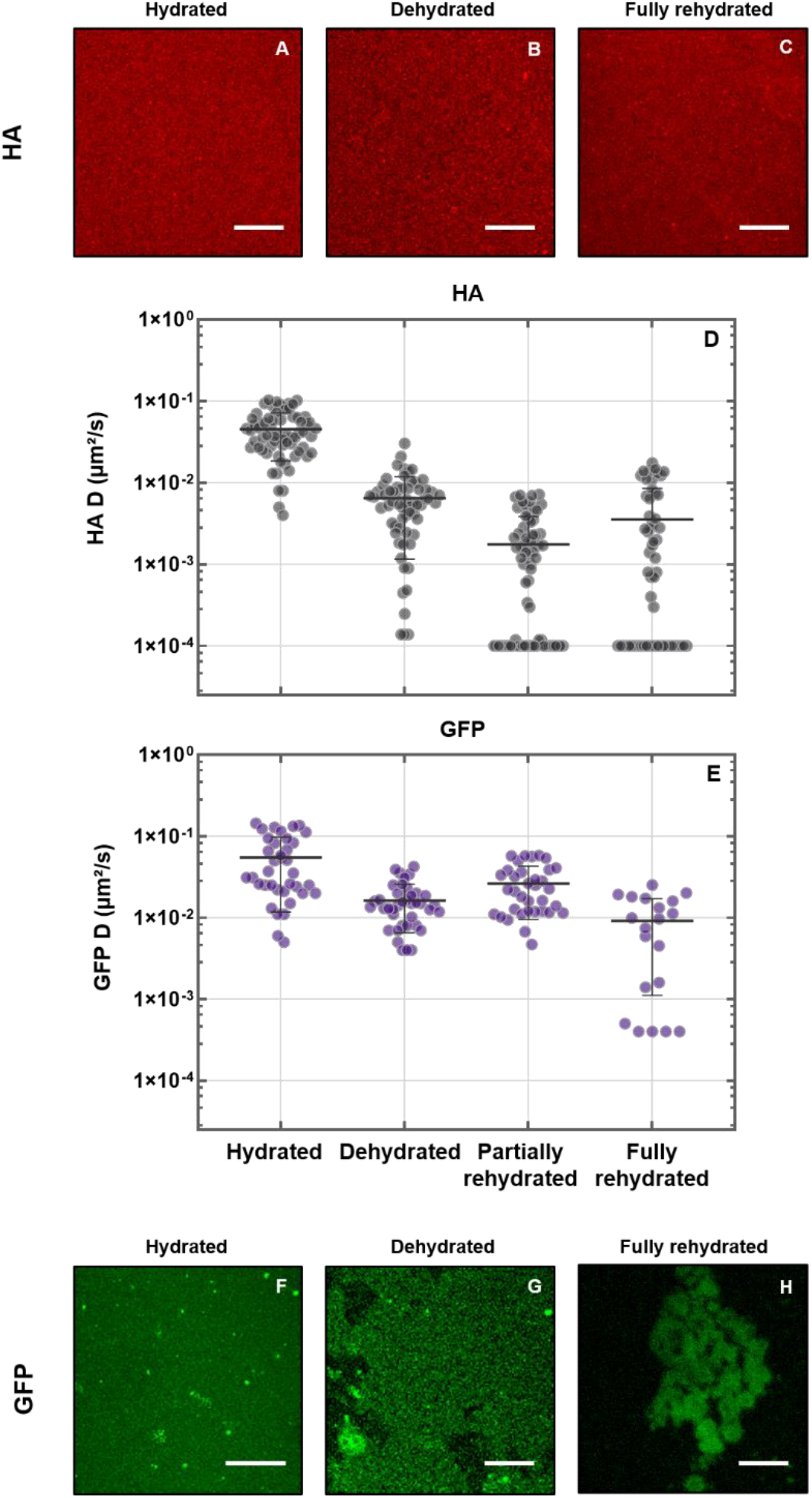
Fluorescence images and diffusion coefficients (D) of proteins in SLBs measured via RICS at different hydration conditions. (A and F) Representative images of proteins in hydrated SLBs immediately after preparation. (B and G) Representative images of proteins in dehydrated SLBs, after removal of bulk water and desiccation at 20 % relative humidity. (C and H) Representative images of proteins in fully rehydrated SLBs in DPBS buffer. (A-C) A568 labeled HA conjugated to DOPC 90 % DGS-NTA-Ni 10 % corresponding to the samples shown in Figure 1 (B, E and H), respectively. (F-H) GFP conjugated to DOPC 90 % DGS-NTA-Ni 10 % corresponding to Figure 1 (C, F and I), respectively. Diffusion coefficients measured via RICS are shown for HA (D) or GFP (E) bound to SLBs in different hydration conditions. HA was labeled with A568 and anchored to SLBs containing DOPC 90% DGS-NTA-Ni 10% and labeled with 0.01% TF-PC. GFP was anchored to SLBs containing DOPC 90% DGS-NTA-Ni 10% and labeled with 0.01% Rho-PE. Both proteins were bound to the bilayer via His-tag and DGS-NTA-Ni. Data were obtained in at least three independent bilayer replicates (ca. 9-12 measurements per replicate). Individual points represent a single RICS measurement on one of the replicates. The horizontal lines show the mean D and the vertical lines the standard deviation. For better visualization, D values below 1 × 10^−4^ μm^2^/s were set as constant and equal to this minimum threshold. Scale bars correspond to 20 μm.

As shown in Figure 4 B, the removal of bulk water following dehydration did not affect macroscopically the HA layer on the SLBs (Figure 4 B). On the contrary, the GFP protein layer was notably damaged (Figure 4 G). In terms of dynamics, sample dehydration caused an almost irreversible decrease in protein diffusion, both for HA and GFP (Figure 4 D and E “dehydrated”). It is worth noting, though, that much slower dynamics were observed in dehydrated or partially rehydrated samples, for the case of HA-bilayers. In the dehydrated state, HA diffusion decreased in fact to ~ 15 % ((0.006 ± 0.005) μm^2^/s) while GFP diffusion decreased to ~ 30 % ((0.016 ± 0.010) μm^2^/s) of the initial values. Fluorescence images of proteins in partially rehydrated samples were comparable to those obtained in dehydrated condition, as for the case of the lipid bilayers described in 3.1 (data not shown). Also, partial rehydration led to a further decrease in HA mobility ((0.002 ± 0.002) μm^2^/s, Figure 4 D “partially rehydrated”) but not for GFP ((0.026 ± 0.017) μm^2^/s, Figure 4 E “partially rehydrated”). Following full rehydration, HA-bilayers remained macroscopically unaltered (Figure 4 C) while GFP-bilayers remained spatially inhomogeneous (Figure 4 H). In these conditions, we obtained D values of (0.003 ± 0.005) μm^2^/s for HA (Figure 4 D “fully rehydrated”) and (0.009 ± 0.008) μm^2^/s for GFP (Figure 4 E “fully rehydrated”).

### 3.3 LSFCS indicates irreversible decrease of HA dynamics and partial recovery of lipid diffusion upon dehydration-rehydration cycles

RICS measurements were performed within a single frame acquisition (ca. 1 s), due to the significant photobleaching observed especially after sample dehydration. Hence, information on protein dynamics could only be obtained on a microsecond to millisecond time scales. Furthermore, since RICS relies on a two-dimensional spatial correlation on a larger region of the sample (compared to LSFCS which consists of a one-dimensional spatial correlation analysis on a selected and restricted region of the bilayer), this approach can be more easily influenced by the presence of slow-moving large fluorescent objects. Therefore, these experiments were complemented by LSFCS measurements in order to obtain information on longer times scales (i.e. several seconds) and, thus, increase the statistical accuracy of the results in the case of very slow dynamics. The diffusion coefficients D of lipids determined via LSFCS were in all cases slightly higher (Figure 5 A, Table S1) compared to those obtained via RICS, as expected. Nevertheless, the general decrease of lipid dynamics upon dehydration did not differ, independently of the employed technique. In the case of fully rehydrated samples, a slight but significant (p < 0.003) increase in lipid dynamics could be observed compared to partially rehydrated samples at variance with the results obtained by RICS (p > 0.20, Figure 2 A).

**Figure 5.**
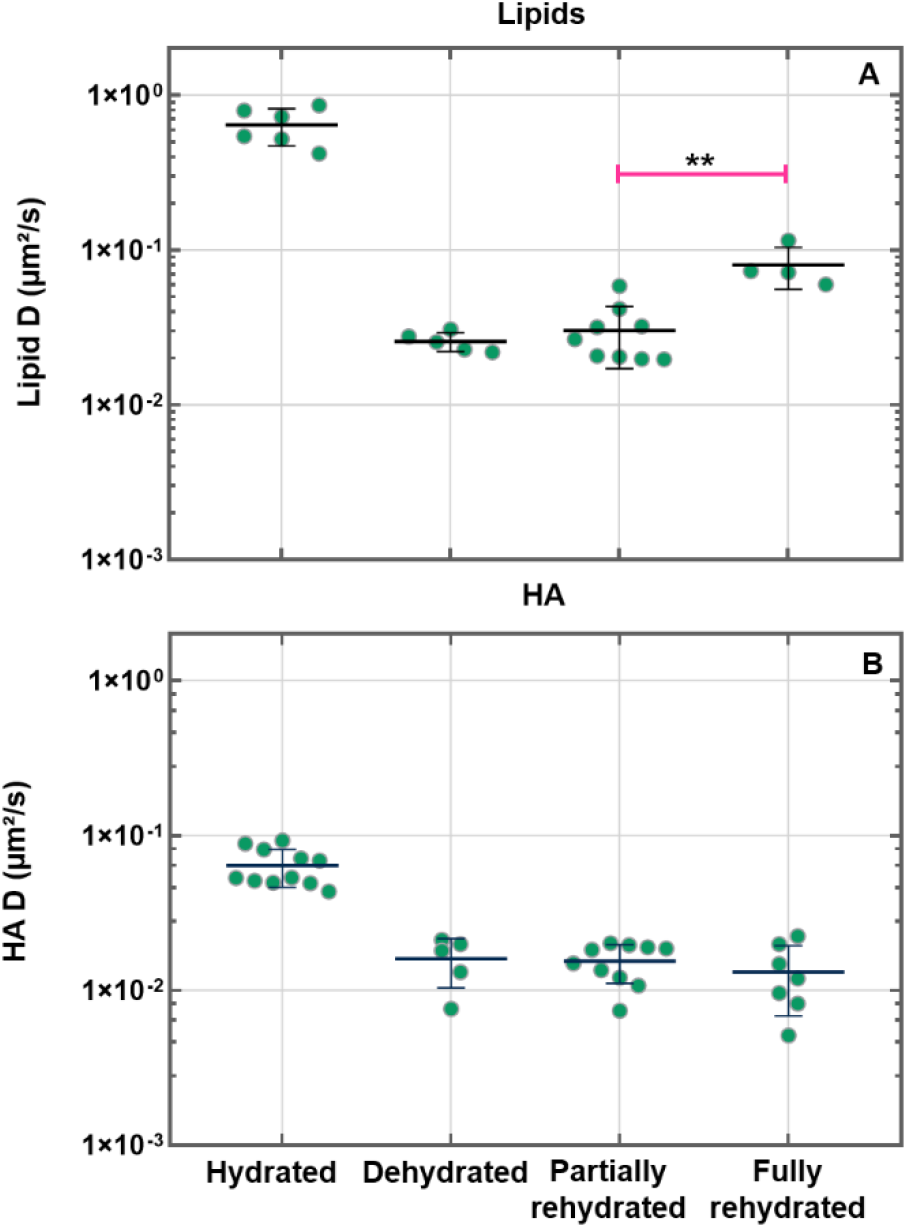
Diffusion coefficients (D) of lipids and HA proteins in SLBs measured via LSFCS at different hydration conditions. HA was labeled with A568 and bound to SLBs containing DOPC 90 % DGS-NTA-Ni 10 % and labeled with 0.01 % TF-PC. HA proteins were anchored to the bilayer via His-tag and DGS-NTA-Ni. Diffusion coefficients measured via LSFCS are shown for the lipids (A) and HA proteins (B). Data were obtained in two independent bilayer replicates (ca. 3-6 measurements per replicate). Individual points represent a single LSFCS measurement on one of the replicates. The grey horizontal lines show the mean D and the vertical lines the standard deviation. The pink horizontal line in (A) indicates a significant but minor increase in lipid diffusion in the fully hydrated samples with respect to the partially rehydrated condition (**p < 0.003).

Similar observations were performed in the case of protein dynamics (Figure 5 B, Table S2). Although D values determined via LSFCS were consistently higher, both techniques indicated the same trend in the reduction of HA diffusion upon dehydration and subsequent sample rehydration.

### 3.4 AFM reveals dehydration-induced membrane disruption at different space scales

We complemented the fluorescence microscopy experiments with AFM measurements, obtaining topography images and local surface characterization on the hydrated, dehydrated, partially rehydrated and fully rehydrated samples (Figure 6).

**Figure 6.**
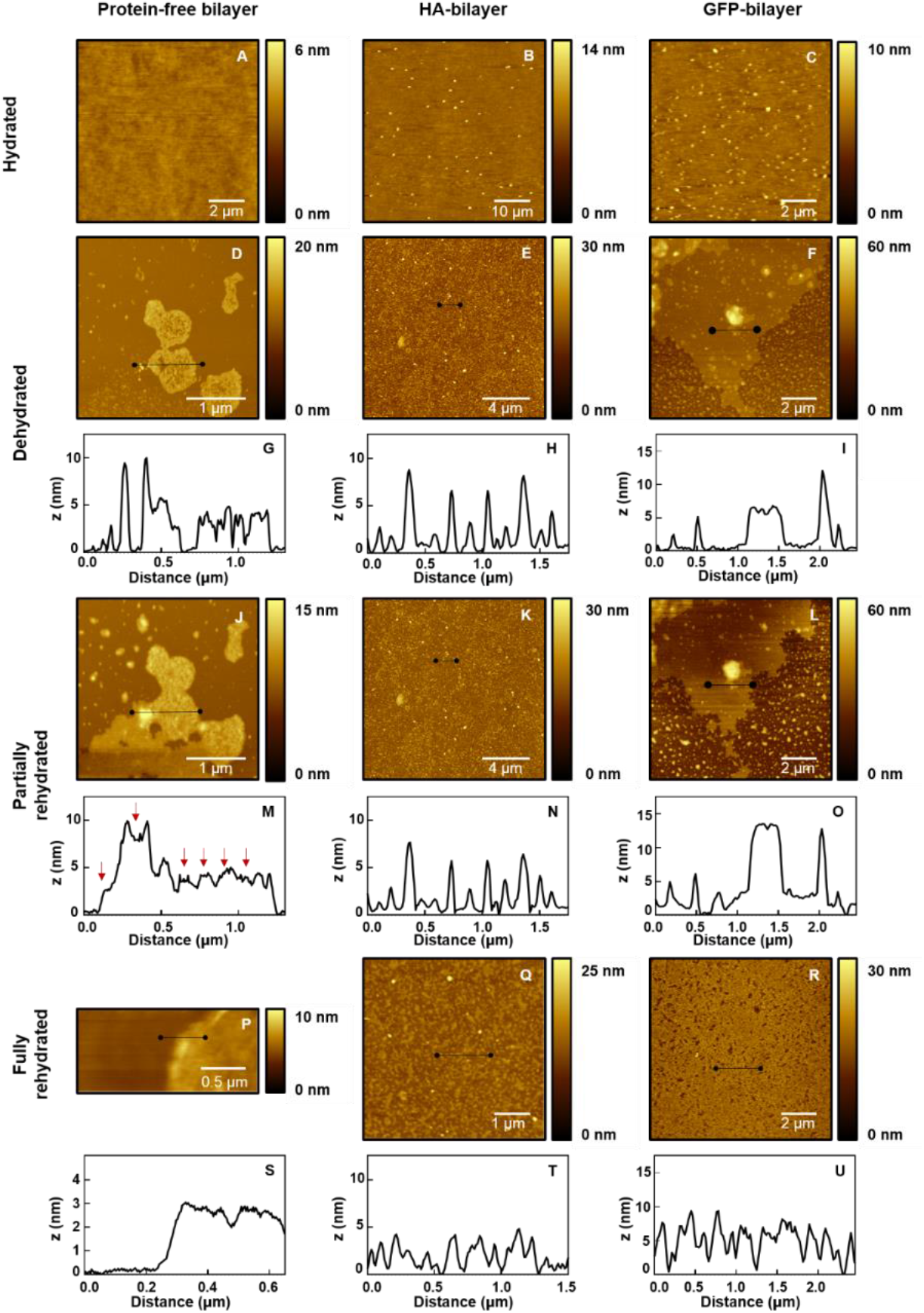
AFM topography images and height profiles of SLBs at different hydration conditions. (A-C) Representative images of hydrated SLBs in DPBS buffer immediately after preparation. (D-I). Representative images of dehydrated SLBs, after removal of bulk water and desiccation at 20 % relative humidity. (J-O) Representative images of partially rehydrated SLBs at 95 % relative humidity. (P-R) Representative images of fully rehydrated SLBs in DPBS buffer. (A, D, J and P) Protein-free bilayers of DOPC 90 % DGS-NTA-Ni 10 %. (B, E, K and Q) Bilayers of DOPC 90 % DGS-NTA-Ni 10 % conjugated to HA 2 μM. (C, F, L and R) Bilayers of DOPC 90 % DGS-NTA-Ni 10 % conjugated to GFP 2 μM. (G and I) Height profiles extracted from the horizontal lines of (D and F), respectively. (M and O) Height profiles extracted from the horizontal lines of (J and L), respectively. (H and N) Height profiles relative to the black lines shown in (E and K), respectively. (S-U) Height profiles extracted from the horizontal lines of (P-R), respectively. Red arrows in (M) point changes in height with respect to (G). HA maximum height in solution was obtained from 8 different images (number of proteins = 104) and for GFP from 5 different images (number of proteins = 98).

Freshly prepared protein-free bilayers appeared homogenous and flat, displaying a surface roughness of (0.43 ± 0.06) nm (Figure 6 A). After anchoring the proteins to the SLBs, we expected to observe high protein density, as detected by fluorescence microscopy. However, probably due to significant tip-sample interactions and the applied force to the soft surface [42] [43] [44], we only observed few proteins anchored to the surface, especially for HA-bilayers. The measured maximum height of the single features observed on protein-bilayers was on average (14.7 ± 2.1) nm for HA (Figure 6 B) and (4.4 ± 1.1) nm for GFP (Figure 6 C). These results agree well with AFM studies on HA [27] [45] and GFP [46], as well as with the crystal structures of these proteins [47] [48].

Following dehydration, we acquired AFM topography images in air. The protein-free bilayers presented major structural damage and extensive areas with uncovered glass (Figure 6 D) and – in agreement with fluorescence microscopy results – the surface coverage decreased to (25 ± 6) %. Specifically, we observed features of diverse sizes: ca. 4 nm high patches (Figure 6 G), structures with height between ~ 4 - 30 nm (Figure S2 A and B) and other objects with height ~ 100 nm (Figure S2 A and C), indicating the probable presence of single bilayers, multiple bilayers and lipid aggregates, respectively.

Topography images on dehydrated HA-bilayers consistently showed homogeneous samples on a macroscopic scale comparable to the fluorescence micrographs (Figure 6 E and H). However, at larger magnification, the high lateral resolution of AFM revealed membrane disruption (Figure S2 D and E). Indeed, we noticed a decrease in the surface coverage down to (61 ± 10) %. Figure 7 A is a typical representative zoom of Figure 6 E and shows homogenous distribution of nanoscale patches on the substrate of ~ 100 - 500 nm lateral size and ~ 4 nm height. The AFM image height histograms (Figure S3) can be fitted to two Gaussian distributions, whose peak locations differ by (3.8 ± 0.6) nm, as expected for a single bilayer on a substrate (Figure S4). We also identified features that protrude ~ 13 nm (Figure 7 A and B), which should correspond to single or aggregated HA proteins anchored to the lipid bilayer.

**Figure 7.**
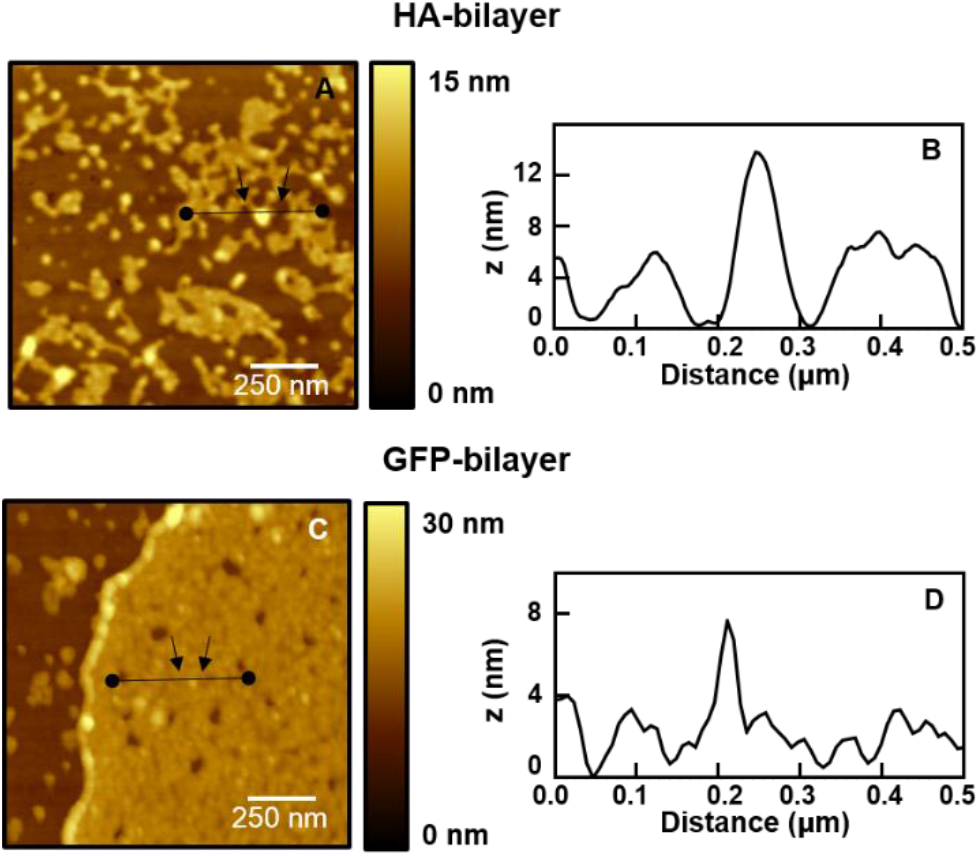
AFM topography images and height profiles of proteins on SLBs after dehydration. (A) Representative image of a SLB of DOPC 90 % DGS-NTA-Ni 10 % conjugated to HA 2μM. (B) Representative image of a SLB of DOPC 90 % DGS-NTA-Ni 10% conjugated to GFP 2 μM. (B and D) Height profiles extracted from the horizontal lines of (A and C), respectively. Black arrows indicate features that could be identified as proteins.

Different from HA-bilayers, but similar to the protein-free bilayers, AFM scans on the microscale showed GFP-bilayers with extensive damage of the bilayer and exposure of the glass surface (Figure 6 F and I, Figure S2 F and G). The surface coverage was of (44 ± 12) %. We observed residual lipids on the glass surface, bilayer patches and lipid aggregates. In some areas we found features of ~ 6 nm that could correspond GFP molecules (Figure 7 C and D).

For the analysis of partially rehydrated samples, we scanned and zoomed in the same areas as for dehydrated bilayers and compared the surface topography between both conditions. On the protein-free bilayers (Figure 6 J), we observed lateral expansion of the remaining lipid patches, i.e. an increase in the surface coverage of ~ 15 %, as shown in Figure 6 D and J. Also, a large number of holes in the SLB, which we assign to regions of exposed solid support, were filled, as observed by comparing the height profiles from Figure 6 G and M (see arrows). The HA-bilayers did not show significant structural alterations upon exposure to high humidity (Figure 6 K and N) while GFP-bilayers revealed membrane rearrangement in shape and height (Figure 6 L and O). For example, we observed topographic features on top of the lipid bilayer that changed in height and shape, upon partial rehydration (Profiles I and O). Such features can be identified as e.g., lipid aggregates swelling upon rehydration.

Finally, we noticed some topographical alterations of the protein-free bilayers and the GFP-bilayers upon full rehydration (Figure 6 P and U). The AFM images in protein-free bilayers revealed non-homogenous bilayer regions (Figure 6 P and S). Similarly, GFP-bilayers were non-uniform with defects and holes (Figure 6 R and U) and large regions of uncovered glass (Figure S5 A and B). Interestingly, fully rehydrated HA-bilayers were comparable to those in condition of dehydration or partial rehydration (Figure 6 Q and T).

## 4. Discussion

In this work we used fluorescence microscopy, RICS, LSFCS and AFM to study the effects of dehydration on the dynamics and structure of models of the influenza virus envelope. We specifically investigated the protective effects of HA at the macro- and microscopic spatial scale from a biophysical point of view.

Our models consisted of PC lipid bilayers with HA incorporated at high concentration to mimic the IAV H1N1 envelope. In hydrated conditions, both membrane structure (Figure 1 A-C, Figure 6 A-C) and lipid dynamics (Figure 2) suggest that these bilayers are in the fluid phase [49] [50] [51]. In bilayers with incorporated proteins, the relatively slow dynamics of lipids (compared e.g. to those reported in [19]) might be ascribed to molecular crowding [52] due to high protein densities (Figure 4).

After sample dehydration, two different phenomena are observed: irreversible macroscopic disruption of the bilayers and microscopic membrane damage. Macroscopic disruption is observed in the scale of tens of micrometers while microscopic membrane damage is observed locally at the nanoscale, in less than 500 nanometer regions. The former phenomenon is hindered specifically by the presence of HA (Figure 1 E and H), as membranes without this protein exhibit clear structural damage (Figure 1 D and F). Such large-scale damage is irreversible and can be still observed also after partial or full rehydration (Figure 1 G and I). In protein-free bilayers, water appears therefore to induce reorganization, desorption, delamination and aggregation of lipid molecules, in line with previous results [18] [53]. Extensive membrane damage also occurs in GFP-bilayers but not in HA-bilayers, suggesting that the protective effect on a macroscopic scale depends on the specific protein identity.

We hypothesize that the HA-induced protective effect is due to the formation of a glycoprotein-water matrix on the membrane when the protein is present at high concentrations. GFP, on the other hand, appears less prone to establish similar intermolecular interactions. HA is a trimeric glycoprotein rich in surface glycans with a high mannose content. It also contains more complex and branched glycans [54]. Surface glycans increase the overall stability of the proteins in solution [55] and mannosyl derivates were shown to support protein integrity upon freeze-drying or desiccation [56] [57]. Furthermore, AFM experiments indicate that mannobiose monolayers present strong self-adhesion forces which, in turn, may play a role in water structuring and retention of a water layer upon bulk dehydration [58]. Finally, exposure of certain organisms to extremely low humidity conditions can induce a so-called “anhydrobiosis” state, a condition in which high contents of trehalose, sucrose or other disaccharides are produced to avoid cell death [59]. While not providing direct evidence regarding the molecular mechanism, our experiments suggest that the high density of surface glycans in HA might be responsible for the formation of a rigid molecular matrix that retains water molecules and allows membrane preservation upon dehydration. A similar effect was previously reported for lipid bilayers which were preserved in dehydrated conditions by a layer of disaccharides [53] or monosaccharides [60]. Analogous findings regarding the stabilization of dehydrated membranes by protein crowding were reported by Holden et al. [17]. In their study, the presence of streptavidin or IgGs at very high density preserved the membrane after exposure to air. It was therefore reported that the protein’s identity may not be determinant for membrane protection. Of interest, both streptavidin and IgGs are glycosylated proteins and the potential influence of the glycan composition on the results was not discussed.

The second type of damage (i.e., microscopic membrane damage) is observed regardless of the membrane composition and the presence of proteins. Water removal induces a phase transition into a gel state [61] and a decrease in the lipid area per headgroup [62], resulting in membrane shrinkage, exposure of the glass substrate and reduction of the total surface coverage of the bilayers (Figure 6 D-F). As a consequence, we observed the formation of microscopic membrane defects which might not be necessarily connected to actual loss of lipids. These small discontinuities in the membrane might further hinder residual lipid diffusion (Figure 2), as also reported before [16]. Protein diffusion decreased as well during dehydration albeit only limitedly, probably due to the already slow dynamics in crowded hydrated samples (Figure 4 D and E).

Upon rehydration, microscopic membrane damage appears reversible in protein-free bilayers and GFP-bilayers. The reintroduction of bulk water favors the rearrangement of lipids on the membranes resulting in membrane expansion over the glass surface [63] and, possibly, a phase transition back to the liquid-crystalline state [64]. In agreement with our quantification of lipid dynamics (Figure 2), the re-establishment of membrane fluidity allows small defects to be repaired and membrane self-healing thus occurs at the micrometric scale (Figure 6 J, L, P and R). This local process does not appear to be influenced by large-scale irreversible macroscopic damage on the bilayer, as reported in analogous investigations [65]. The larger data spread observed for lipid or protein diffusion coefficients in protein-containing SLBs suggests that these samples are spatially inhomogeneous.

On the other hand, HA-bilayers exhibit irreversible microscopic membrane damage. Rehydration does not change the apparent structure of the bilayers (Figure 6 K and Q), as the samples remain comparable to the dehydrated state. Also, lipid and protein dynamics are only marginally recovered (Figure 2, Figure 4, Figure 5). One possible explanation is that a rigid matrix of intermolecular interactions among residual water molecules and surface glycans of HA stabilizes the bilayer is a solid gel-state. Such bilayer phase is induced by dehydration (as also observed in protein-free and GFP bilayers) but is not reversed by rehydration. Of interest, a suppression of gel to liquid-crystalline phase separation during membrane rehydration was associated to the prevention of content leakage from lipid vesicles [64].

In summary, our model of the IAV H1N1 envelope appears to undergo a transition from fluid to solid phase upon dehydration. It worth noting though that IAV lipids are more likely to be found in a solid gel-like state rather than in a fluid disordered phase, already when fully hydrated [66] [67]. This is due to the fact that the viral envelope is highly enriched in phosphatidylethanolamine (PE) and phosphatidylserine (PS) with much lower amounts of unsaturated PC, compared to e.g. the plasma membrane of the host cell [68] [69]. On one hand, the fact that our model envelope is in the fluid phase in hydrated samples allows us to monitor protein and lipids dynamics as a function of water content. On the other hand, this represents a limitation of our system since the observed fluid-to-solid phase transition might not occur in IAV virions upon dehydration. We hypothesize therefore that the irreversible microscopic damage that we see in our samples does not play a major role in the process of virus survival at low humidity. In contrast, our data suggest that HA might play a fundamental role in protecting the virus envelope from large-scale damage and disintegration during dehydration.

## 5. Conclusions

We developed a novel approach to investigate the biophysical properties of the IAV H1N1 envelope under different hydration conditions from the macro- to the microscale. While our model is mainly limited by the specific membrane composition, our results indicate that HA might act as a membrane protectant upon dehydration. This observation provides new insights regarding the molecular mechanisms determining virus stability and survival outside living cells. In general, the experimental approach presented here takes advantage of the synergy between AFM and fluorescence microscopy since the two techniques provide spatial information on different spatial scales. Future studies could address the role of specific lipids and surface glycans in determining the interactions between viral envelope and water molecules.

## 6. Author contributions

**Maiara A. Iriarte-Alonso**

Conceptualization, Formal analysis, Funding acquisition, Methodology, Project administration, Resources, Software, Supervision, Validation, Visualization, Writing - original draft, Writing – review and editing.

**Alexander M. Bittner**

Conceptualization, Formal analysis, Funding acquisition, Methodology, Project administration, Resources, Supervision, Visualization, Writing – review and editing.

**Salvatore Chiantia**

Conceptualization, Formal analysis, Funding acquisition, Methodology, Project administration, Resources, Software, Supervision, Visualization, Writing – review and editing.

## 8. Acknowledgments

The authors thank Dr. Kerstin Blank and Reinhild Dünnebacke (Max Planck Institute of Colloids and Interfaces, Mechano(bio)chemistry Group) for providing with an AFM instrument during M.A.I.A research visit at Potsdam (2020). We thank Aitziber Eleta-Lopez (CIC nanoGUNE (BRTA), Self-Assembly Group), and also Annett Petrich and Zaloa Aguirre Sourrouille (University of Potsdam, Cell Membrane Biophysics Group) for valuable discussions.

This work was supported by the German Research Foundation [DFG grant number 254850309 to S.C]; the Basque Government [grant number Elkartek 2019 to A.M.B]; the Spanish MINECO/MCIU [grant number PID2019 104650GB and Maria de Maeztu “Units of Excellence” Program MDM 2016 0618 to A.M.B]; the Project Cursum ADASTRA [project “Wetinflu TRAV” to A.M.B], the European Molecular Biology Organization [EMBO Short-Term Fellowship grant number 8397 to M.A.I.A] and the German Academic Exchange Service [DAAD, Research Grants-Short-Term Grants 2020 number 57507442 to M.A.I.A].

## Supplementary information

**Figure S1.**
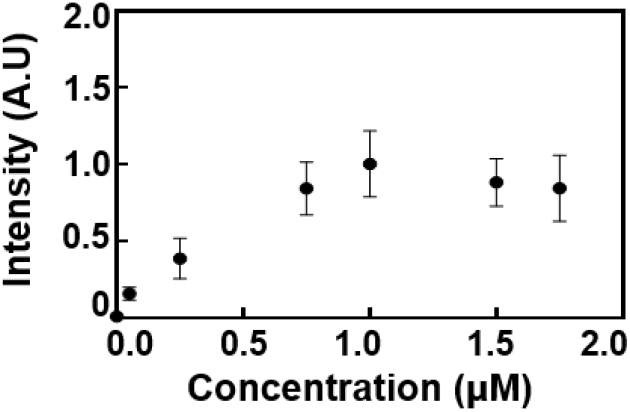
Binding curve of His-tagged HA performed by fluorescence microscopy. Individual points represent the mean intensity at each condition and the vertical line the standard deviation.

**Table S1.**
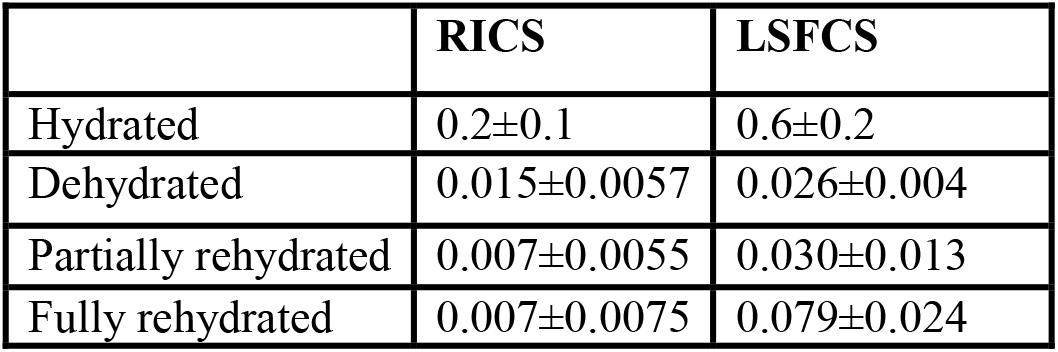
Diffusion coefficients (D) of lipids in HA-bilayers measured via RICS and LSFCS at different hydration conditions. All examined SLBs contained DOPC 90 % DGS-NTA-Ni 10 % and were labeled with 0.01 % TF-PC. HA proteins were labeled with A568 and were anchored to the bilayer via His-tag and DGS-NTA-Ni. RICS data were obtained in at least three independent bilayer replicates (ca. 9-12 measurements per replicate) while LFCS data were obtained in two independent bilayer replicates (ca. 3-6 measurements per replicate).

**Table S2.** Diffusion coefficients (D) of HA proteins in HA-bilayers measured via RICS and LSFCS at different hydration conditions. All examined HA were labeled with A568 and conjugated to SLBs containing DOPC 90 % DGS-NTA-Ni 10 % and labeled with 0.01 % TF-PC. HA proteins were anchored to the bilayer via His-tag and DGS-NTA-Ni. RICS data were obtained in at least three independent bilayer replicates (ca. 9-12 measurements per replicate) while LFCS data were obtained in two independent bilayer replicates (ca. 3-6 measurements per replicate).

|  | RICS | LSFCS |
| --- | --- | --- |
| Hydrated | 0.045±0.0265 | 0.064±0.0175 |
| Dehydrated | 0.0065±0.0053 | 0.0159±0.0056 |
| Partially rehydrated | 0.0018±0.0021 | 0.0154±0.0044 |
| Fully rehydrated | 0.0035±0.0050 | 0.0131±0.0063 |

**Figure S2.**
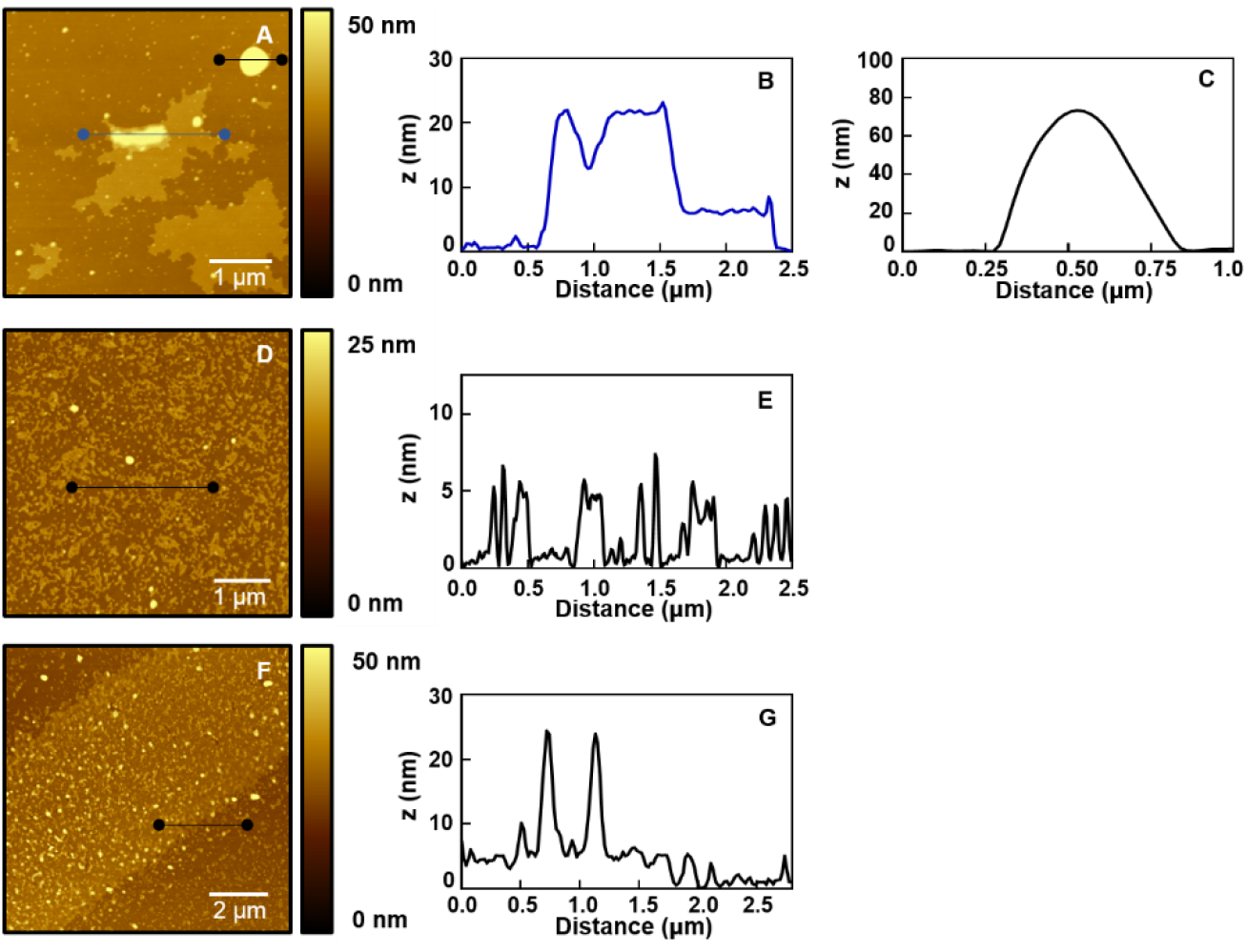
AFM topography images and height profiles of SLBs after dehydration. (A) Representative image of protein-free bilayers of DOPC 90 % DGS-NTA-Ni 10 %. (B and C) Height profiles extracted from the blue and black horizontal lines in (A), respectively. (D) Representative image of bilayers of DOPC 90 % DGS-NTA-Ni 10 % conjugated to HA 2 μM. (E) Height profile extracted from the horizontal line in (D). (F) Representative image of bilayers of DOPC 90 % DGS-NTA-Ni 10 % conjugated to GFP 2 μM. (G) Height profile extracted from the horizontal line in (F).

**Figure S3.**
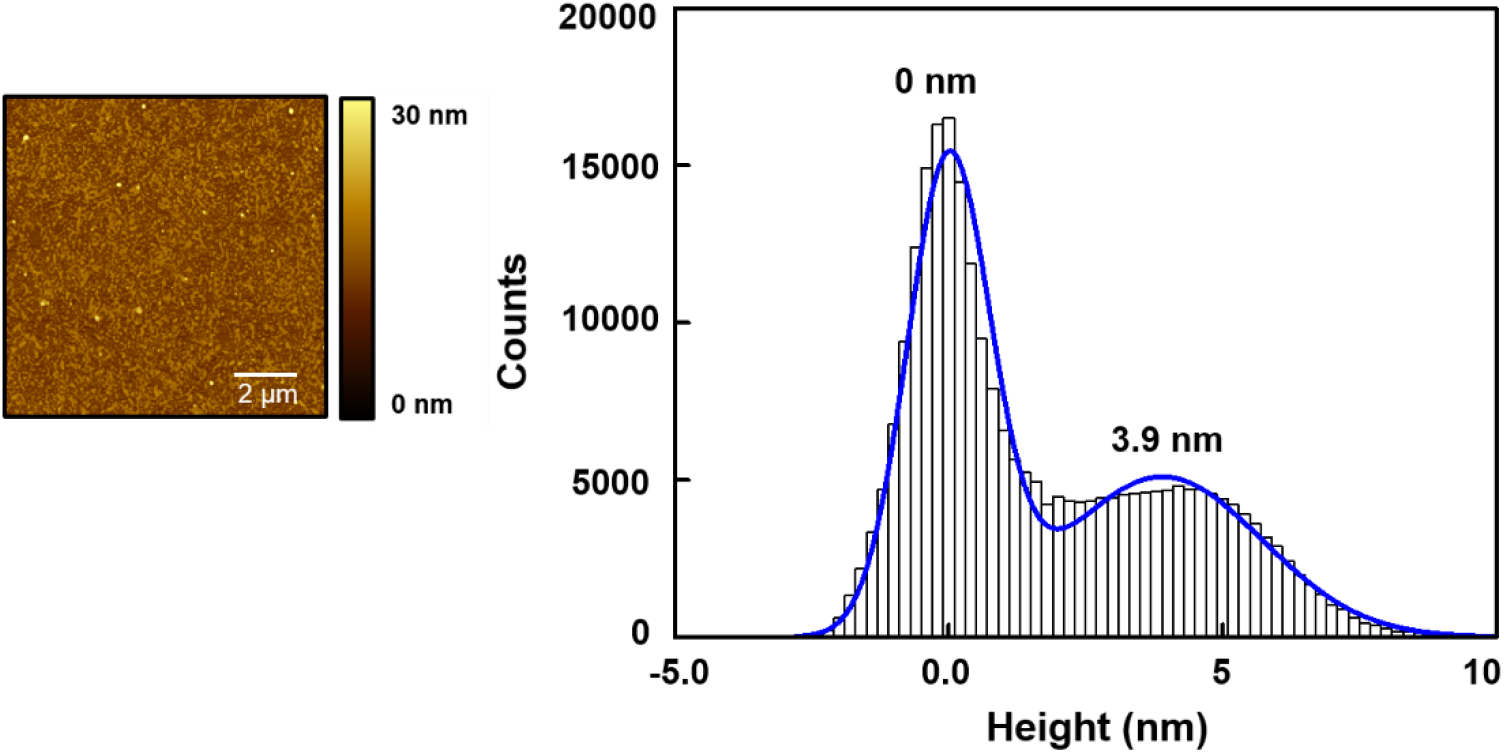
AFM topography image and height histogram of HA-bilayers after dehydration. The height histogram was fitted to a 2-peak Gaussian function (number of images, n = 16). The estimated height difference from the main peaks was considered as the depth of the lipid bilayer in the dehydrated state. Note that the height data provided by the AFM are not absolute values. Hence, we assigned arbitrarily the average height of the substrate underneath the bilayer to ~ 0 nm. Also, the Gaussian fit is not required to obtain the result, but it makes sense in terms of assigning statistical height variations.

**Figure S4.**
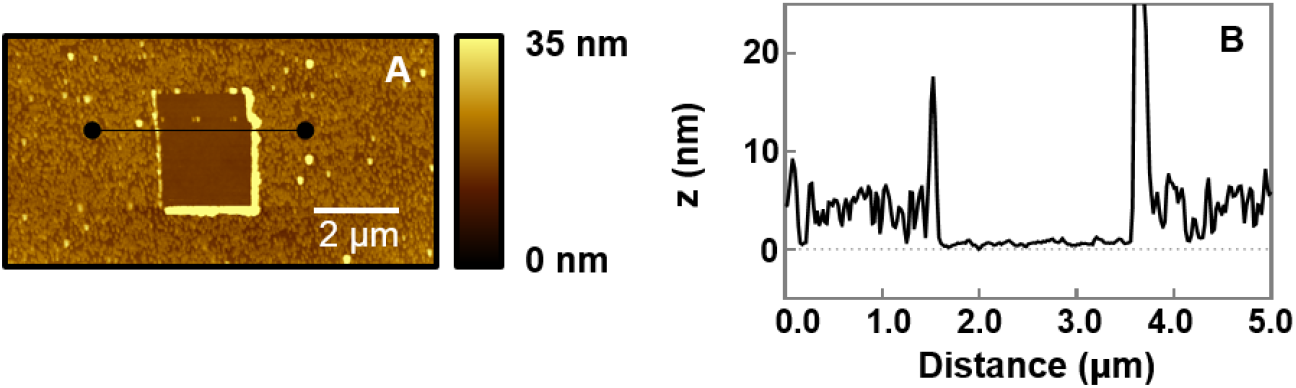
AFM topography image and height profile of HA-bilayers obtained from the scratch and scan test after dehydration. (A) Representative AFM topography image of a scratched region on DOPC 90% DGS-NTA-Ni 10% SLB containing HA. (B) Height profile extracted from the horizontal line in (A).

**Figure S5.**
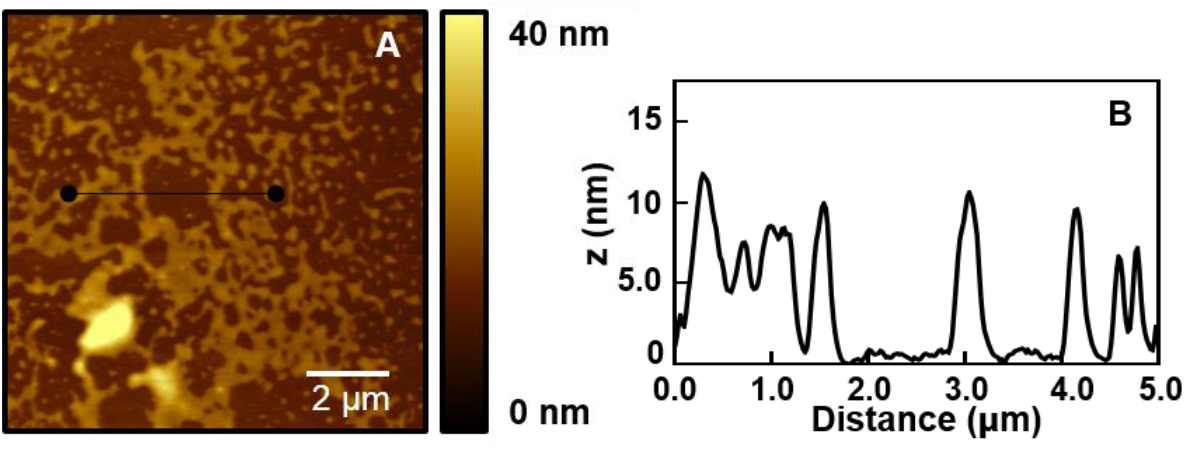
AFM topography image and height profile of GFP-bilayers after full rehydration. (A) Representative AFM topography image of DOPC 90% DGS-NTA-Ni 10% SLB containing GFP. (B) Height profile extracted from the horizontal line in (A).

